# SAILER: Scalable and Accurate Invariant Representation Learning for Single-Cell ATAC-Seq Processing and Integration

**DOI:** 10.1101/2021.01.28.428689

**Authors:** Yingxin Cao, Laiyi Fu, Jie Wu, Qinke Peng, Qing Nie, Jing Zhang, Xiaohui Xie

**Affiliations:** Department of Computer Science, University of California, Irvine, CA, 92697, USA; Systems Engineering Institute, School of Electronic and Information Engineering, Xi’an Jiaotong University, Xi’an, Shannxi, 710049, China; Department of Biological Chemistry, University of California, Irvine, CA, 92697, US; Department of Mathematics, University of California, Irvine, CA, 92697, USA; Center for Complex Biological Systems, University of California, Irvine, CA, 92697, USA; NSF-Simons Center for Multiscale Cell Fate Research, University of California, Irvine, CA, 92697, USA.

## Abstract

**Motivation:** Single-cell sequencing assay for transposase-accessible chromatin (scATAC-seq) provides new opportunities to dissect epigenomic heterogeneity and elucidate transcriptional regulatory mechanisms. However, computational modelling of scATAC-seq data is challenging due to its high dimension, extreme sparsity, complex dependencies, and high sensitivity to confounding factors from various sources.

**Results:** Here we propose a new deep generative model framework, named SAILER, for analysing scATAC-seq data. SAILER aims to learn a low-dimensional nonlinear latent representation of each cell that defines its intrinsic chromatin state, invariant to extrinsic confounding factors like read depth and batch effects. SAILER adopts the conventional encoder-decoder framework to learn the latent representation but imposes additional constraints to ensure the independence of the learned representations from the confounding factors. Experimental results on both simulated and real scATAC-seq datasets demonstrate that SAILER learns better and biologically more meaningful representations of cells than other methods. Its noise-free cell embeddings bring in significant benefits in downstream analyses: Clustering and imputation based on SAILER result in 6.9% and 18.5% improvements over existing methods, respectively. Moreover, because no matrix factorization is involved, SAILER can easily scale to process millions of cells. We implemented SAILER into a software package, freely available to all for large-scale scATAC-seq data analysis.

**Availability:** The software is publicly available at https://github.com/uci-cbcl/SAILER

## 1 Introduction

Accessible chromatin regions host a network of complex interplays among numerous cis-regulatory elements (CREs, such as enhancers and promoters), transcription factors (TFs), cofactors, and chromatin remodelers in the three-dimensional genome for precise spatiotemporal gene expression control (Klemm *et al*., 2019; Tsompana and Buck, 2014; Boyle *et al*., 2008). Assay for transposase-accessible chromatin using sequencing (ATAC-seq) is an efficient method to probe accessible DNA regions in the genome, by tagging them with sequencing adapters using the Tn5 transposase (Buenrostro, Wu, Chang, *et al*., 2015). More recently, researchers have developed single-cell ATAC-seq (scATAC-seq) technology to massively probe accessible chromatin regions in individual cells (Buenrostro, Wu, Litzenburger, *et al*., 2015; Cusanovich *et al*., 2015; Chen *et al*., 2018; Satpathy *et al*., 2019). These methods make it possible to comprehensively dissect the epigenetic heterogeneity across diverse cell states at an unprecedented resolution. Due to its easy protocols and high-throughput capacities, many labs and big consortia (e.g., the Human Cell Atlas, Human BioMolecular Atlas Program) have employed scATAC-seq for single-cell epigenetic profiling (Regev *et al*., 2017; Consortium and others, 2019). Furthermore, the scientific community and funding agencies have initiated essential data-sharing policies for expedited translational research. Thus, there is an urgent and essential need to develop robust, accurate, and scalable computational methods for scATAC-seq data analysis and integration at a large scale.

Unfortunately, computational modeling of scATAC-seq data has faced several challenges. First, scATAC-seq data tends to have very low coverage, usually with a few thousand distinct reads representing hundreds of thousands to even millions of accessible regions. Second, scATAC-seq contains a high degree of dependencies because numerous cell-type-specific CREs in accessible chromatin regions work in concert to jointly decide cell fate. Lastly, scATAC-seq analysis is highly sensitive to numerous confounding factors arising within and across samples (e.g., read depth variation and dataset-specific conditions).

Researchers have developed many computational approaches to tackle high-dimensional and sparse scATAC-seq data (Schep *et al*., 2017; Fang *et al*., 2019; González-Blas *et al*., 2019; Xiong *et al*., 2019; Fu *et al*., 2020), but each has its limitations. For instance, ChromVAR ignores the impacts of individual peaks and only groups cells by the TF motif enrichment scores from all peaks, resulting in non-optimal clustering performance (Schep *et al*., 2017). SnapATAC uses Jaccard distance to calculate cell-to-cell similarities for dimension reduction with a hidden assumption that peaks are independent of each other and contribute equally to the similarity measure, which is incorrect in most cases. More recently, researchers developed the latent semantic index (LSI) for learning the lower-dimensional cell representations (Pliner *et al*., 2018; Granja *et al*., 2020; Stuart *et al*., 2020). Despite their scalability, such linear techniques may not fully capture the complex dependencies of peaks. Moreover, these approaches correct for read depth effects by removing components that highly correlate with the read depth, which is heuristic and may lose the true cell-state-related information. Other nonlinear approaches, such as cisTopic and SCALE, were then developed to learn better cell representations (González-Blas *et al*., 2019; Xiong *et al*., 2019). However, these methods assume constant read depths across different cells and ignore potential batch effects from multiple samples, which compromises model performance in real applications.

Here, we aimed to overcome the limitations of existing methods by designing an invariant representation learning scheme with a straightforward intuition – the true epigenetic variations from a specific cell state should remain the same across cells and samples, while variations arising from confounding factors may change substantially, even for cells within similar biological groups. In other words, we can dissect the scATAC-seq cell-to-cell variations into an invariant component representing its hidden cell states and a varying component due to non-biological factors, such as the number of fragments in a cell and batch effects in the multi-sample analyses (**Fig. 1**). To this end, we developed a <u>*s*</u>calable and <u>*a*</u>ccurate <u>*i*</u>nvariant representation <u>*le*</u>a<u>*r*</u>ning scheme (SAILER) via a deep generative model to learn a robust cell representation ***z*** that is only related to intrinsic cell states but is invariant to changes in the confounding factor ***c*** (**Fig. 1**). Specifically, we remove the variations related to confounding factors from the learned latent representation by minimizing their mutual information *I*(***z***, ***c***). Compared with previous methods, SAILER has **three major advantages**: i) it is easily scalable to millions of cells in large-scale analyses via accelerated computation on graphic processing units (GPUs); ii) it captures the nonlinear dependencies among peaks via the expressiveness of deep generative modeling and robustly removes confounding factors from various sources, both within and across samples, to faithfully extract biologically relevant information; iii) it provides a unified strategy for scATAC-seq denoising, clustering, and imputation.

**Fig. 1.**
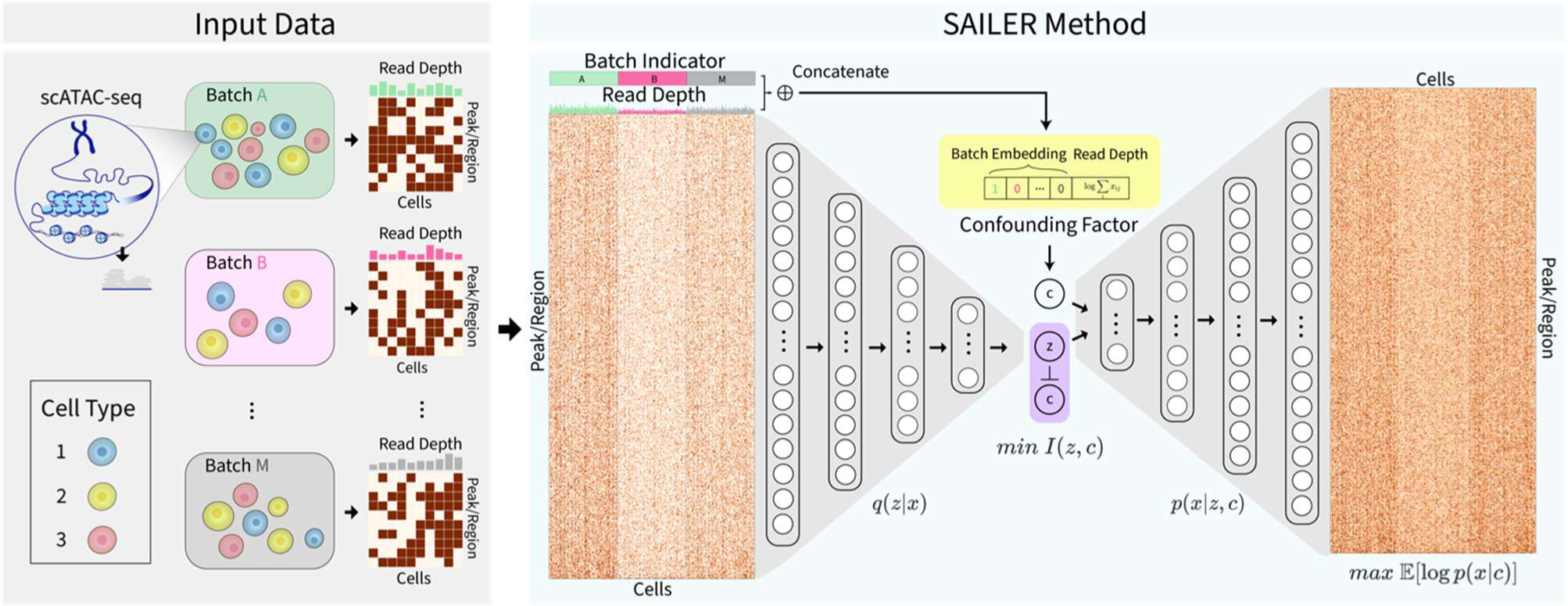
The overall design of the SAILER method. SAILER takes scATAC-seq data from multiple batches as input. Raw data is pushed through the encoder network to obtain a latent representation. Confounding factors for each single cell are concatenated and fed to the decoder along with the latent representation. Batch information is indicated by a one-hot embedding, and read depth is subject to log transform and standard normalization. To learn a latent representation invariant to changes in confounding factors, mutual information between the latent variables and confounding factors are minimized during training.

We implemented SAILER into a Python package that is freely available to the community. To prove its effectiveness, we first benchmarked the clustering performance of SAILER with *state-of-the-art methods*. We utilized three simulated scATAC-seq datasets with ground-truth labels, representing different application scenarios with single- and multi-sample inputs. SAILER significantly outperformed the existing methods, providing improved cell clustering results and successfully identifying rare cell types. We also applied SAILER on real atlas-level and multi-sample scATAC-seq datasets and showed that it could efficiently learn better biologically relevant cell latent representations, which will facilitate various downstream analyses such as cell clustering and imputations.

## 2 Methods

In this section, we provide the mathematical details on our SAILER model and describe methods for benchmarking with existing methods using both simulated and real datasets.

### 2.1 Effective invariant representation learning via a deep generative model

Let ***x*** ∈ {0,1}^*n*^ (with *n* peaks or bins) denote the genome-wide chromatin profile of a cell, with *x*_*i*_ indicating the presence or absence of a peak in bin i. ***x*** depends on both the intrinsic properties of the cell and experimental confounding factors. Our goal is to derive a latent representation of ***x*** (also called embedding) for each cell that reflects only its intrinsic properties. Let ***z*** ∈ *R*^*d*^ be such a latent representation. Suppose ***c*** is the confounding variable that has statistical dependence on ***x***, and is observable together with ***x***. We denote *q* _θ_(***z***|***x***) as the encoder probability, *p*_*ϕ*_ (***x***|***z***, ***c***) as the decoder probability. The decoder part of our model aims to model the conditional probability of ***x*** on ***c*** through a latent variable ***z***,

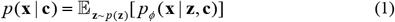

where *p*(***z***) is the prior distribution for a generative model set to be 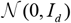 (factorized Gaussian) in our case. *q*(***x***, ***c***) is the empirical distribution of the data point and confounding variable, ϕ denote the parameters of the decoder network.

Following the variational autoencoder (VAE) model (Kingma and Welling, 2014), we performed parameter inference by maximizing an evidence lower bound of the log likelihood, corresponding to minimizing the following loss function,

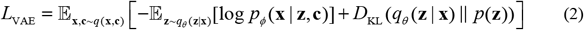

where *q* _θ_(***z***|***x***) is the posterior distribution modeled with a neural net with parameters θ.

The distribution of the latent representation ***z*** induced by empirical data distribution **x** ~ *q*(**x**) = ∑_*c*_*q*(**x, c**) and the posterior probability *q* _θ_(***z***|***x***) potentially can depend on ***c***, as ***c*** is involved in the data generation process. To derive a latent representation ***z*** independent of the confounding variable ***c***, we added an additional term to the loss function to minimize the mutual information between the two variables (Moyer *et al*., 2018),

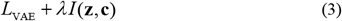

where *I*(***z***, ***c***) is the mutual information between latent representation ***z*** and ***c***, with their joint distribution represented by *q*_θ_(***z***, ***x***, ***c***) = *q*(***x***, ***c***)*q*_θ_(***z***|***x***). Based on the properties of mutual information and variational inequality, *I*(***z***, ***c***) is upper bounded by

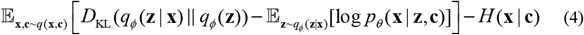

where the conditional entropy *H*(***x***|***c***) is a constant and can be removed from the loss function.

The final loss function we aimed to minimize is

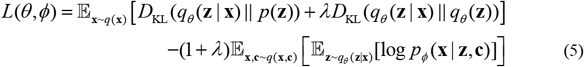

Here D_KL_(*q*_ϕ_ (**z** | **x**) || *p*(**z**)) is the KL-divergence between the encoder *q* _θ_(***z***|***x***) and prior *p*(***z***). 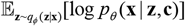 is the reconstruction loss. D_KL_(*q*_ϕ_ (**z** | **x**) || *q*_*θ*_(**z**)) is the KL-divergence between *q* _θ_(***z***|***x***) and empirical marginal distribution *q* _θ_(***z***). Because *q* _ϕ_(***z***) depends on the distribution of both x and c, minimizing the above KL-divergence will reduce the effect of c on z. In the implementation, this extra term is approximated by pairwise KL-divergences between all data points in a training batch,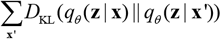. Since latent variable ***z*** is parameterized by an isotropic Gaussian, the pairwise KL has a nice analytical form, and can be efficiently computed with matrix algebra.

### 2.2 Model architecture and training

Considering the close to binary nature of scATAC-seq data, we use binomial likelihood to parameterize the reconstruction loss. To tackle the extreme sparsity issue, we add a positive weight ω to non-zero entries of binary cross-entropy loss 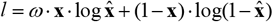 with ω determined by the empirical 0/1 ratio of the input data.

The encoder and decoder are parameterized by two symmetric fully connected feedforward neural networks (with 1000-100-10 units). A sigmoid activation is used for the final output layer. For confounding factors, we use one-hot batch embedding and normalized log-transformed sequencing depth for each cell. During training, input data is pushed through the encoder network to generate the latent variable. Confounding factors are then concatenated together with latent variables and fed into the conditional decoder for reconstruction. As suggested in (Fu *et al*., 2019), when training our model, we adopt a deterministic warmup and cyclical annealing schedule to tackle the KL vanishing problem. Adam optimizer (Kingma and Ba, 2017) with weight decay 5e-4 and minibatch training are used to optimize the model. The model is built with PyTorch library (Paszke *et al*., 2019).

### 2.3 Dimension reduction and clustering

We project the raw high-dimensional sparse scATAC-seq data to a low-dimensional space that reflects the hidden cell states rather than noise in the sequencing experiment. Specifically, we used the raw scATAC-seq matrix ***x*** as the input to our SAILER encoder and extracted the mean of the invariant component ***z*** as the cell representation. We set the default dimension *d* for ***z*** to 10 in our analysis. We then acquired 2D visualizations by running t-distributed stochastic neighbor embedding (t-SNE) (Maaten and Hinton, 2008) or uniform manifold approximation and projection (UMAP) (McInnes *et al*., 2018) on the latent mean. We further constructed a k-nearest neighbor (KNN) graph from the lower-dimensional representations, and then applied the Louvain algorithm (Blondel *et al*., 2008) to assign cells to different clusters.

### 2.4 scATAC-seq imputation

We generated the imputation data via a reconstruction conditioned on the invariant representation z and fixed confounding factor c. Specifically, we first pushed the raw data through the encoder network, and obtained the mean parameters for z. Unlike the training process, where we calculated the depth of the raw data and loaded the one-hot embedding according to the real batch information, here we fixed the depth and batch indicator as the mean depth and the indicator of the batch with the highest data quality. Finally, we concatenated the fixed confounding values with the latent representation z and fed them into the conditional decoder to obtain the imputed data. As a result, we used only the invariant component ***z*** to reconstruct the chromatin landscape during the imputation process, while keeping the other confounding factors at a fixed level.

### 2.5 Performance benchmarking using multiple simulated datasets

We applied SAILER on three simulated scATAC-seq datasets with known cell type labels generated by SCAN-ATAC-Sim (Chen *et al*., 2020) to represent three major application scenarios. We used the peripheral blood mononuclear cell bulk ATAC-seq dataset provided on the SCAN-ATAC-Sim website using all default parameter settings. Each simulation includes three major parameters: *ρ* represents the signal-to-noise ratio (percentage of reads in the true peak regions); *μ* and *σ* denote the mean and standard deviation of the fragment count per cell, respectively. SCAN-ATAC-Sim randomly selects read counts for each cell from a log-normal distribution, and then samples reads from both peak and background regions accordingly. We first simulated a deeply sequenced scATAC-seq dataset (Sim1) with 5,000 fragments per cell (*μ* = 5,000, *σ* = 1.5, and *ρ* = 0.4), representing a scenario in which we are looking for rare cell types. Specifically, we generated 10,000 cells from five cell types, with 100 cells from a rare cell type accounting for 1% of the total population. Then, we generated one shallowly sequenced sample with nine cell types, with *μ* = 3,000, *σ* = 1.5, and *ρ* = 0.4 (Sim2). Lastly, we simulated a two-sample dataset with slightly mismatched cell types to represent scATAC-seq data integration applications with noticeable batch effects – one shallowly sequenced sample (*μ* = 2,500) along with another deep-sequenced sample (*μ* = 5,000) with different signal-to-noise ratios (*ρ* = 0.4 and 0.5, respectively) (Sim3). In addition, we introduced one sample-specific rare cell type in Sim3 to mimic a situation in which rare cell types (e.g., tumor cells) may only exist in some samples. We benchmarked SAILER’s clustering performance with the linear dimension reduction method LSI and another deep learning method, SCALE, on all three simulated datasets. Specifically, we projected the raw input matrix *x* to a ten-dimensional latent space, and further used UMAP to reduce the dimension to 2 for 2D visualization of the cell state landscape. We plotted colored labels according to the ground-truth cell type for visual inspection of clustering performance.

We also used the mutual information to quantify the impacts of confounding factors on the lower-dimensional representations learned by different methods. Specifically, we used a non-parametric mutual information estimation approach (Kraskov *et al*., 2004) to estimate the mutual information between the confounding factors and each dimension of the latent representation, and calculated their mean values for comparison.

### 2.6 Imputation performance on simulated datasets

We also benchmarked the imputation performance of SAILER against SCALE (Xiong *et al*., 2019) and MAGIC (van Dijk *et al*., 2018) on the Sim3 dataset. SCALE is the only current method designated for imputing scATAC-seq data, and MAGIC, originally designed to impute scRNA-seq data, has been incorporated into many scATAC-seq computational pipelines (Fang *et al*., 2019; Granja *et al*., 2020) for imputation purposes.

For SCALE, we directly used the binary imputation output generated by thresholding at mean values of each row and column. For MAGIC, we followed the standard pipeline by applying the recommended *l*1 normalization and square root transformation before imputing the data. Due to the extreme dimension, we used an approximate solver for efficiency. For SAILER, we performed imputation as described in 2.3.

To evaluate the result quantitatively, we calculated the Dice similarity coefficient (DSC) of imputed data 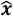 generated by the three methods against the bulk ATAC-seq data ***x***_*bulk*_ of the corresponding cell type used to generate the simulated data. We calculated the DSC of the raw input against the bulk data to provide a baseline.

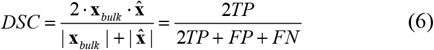

We also generated a 2D visualization to evaluate the landscape of the imputed data. We directly applied a randomized principal component analysis (PCA) (Halko *et al*., 2011) to the imputed data, and used UMAP to visualize the top ten principal components. We also provided the raw input as a baseline.

### 2.7 Performance benchmarking on the mouse atlas dataset

We then demonstrated the performance of our method on a mouse atlas dataset containing 81,173 adult mouse cells from 13 tissues and 40 cell types (Cusanovich *et al*., 2018). Each cell type is annotated by borrowing label information, inferred by marker genes, from the RNA-seq data. A previous effort applied the mouse atlas dataset to benchmark multiple computational methods on scATAC-seq data (Chen *et al*., 2019). The leading method in that study, SnapATAC, was the only method that could process the entire mouse atlas dataset within a reasonable time (~12 h). Given that both SAILER and SCALE are deep learning methods that can train and evaluate data in mini batches, they are capable of handling the scale of the mouse atlas dataset. Thus, we benchmarked SAILER against SnapATAC and SCALE on this dataset.

For SCALE and SAILER, we added a filtering process before loading the data. The filtering involved reducing the bin numbers according to the procedure for filtering peaks used in SCALE. For each cell, we removed bins with read counts of over 90% cells and less than 1% cells.

We used normalized mutual information (NMI) and the adjusted Rand index (ARI) to compare each method’s clustering results with the given labels.

For clustering, we constructed a KNN graph and applied the Louvain algorithm (Blondel *et al*., 2008) to assign clusters to each cell. We compared the clustering results with ground-truth labels to generate the ARI and NMI metrics. We also calculated mutual information between latent representation and confounding factors for comparison.

### 2.8 Performance benchmarking on multi-sample scATAC-seq datasets for mouse brain

To evaluate the ability of SAILER to deal with batch effects, we combined two mouse brain datasets: a mouse brain dataset from the 10X Genomics website and a mouse secondary motor cortex dataset (i.e., the MOs-M1 dataset) (Fang *et al*., 2019). We first selected cells based on barcodes from the 10X mouse brain dataset. Then, we set a threshold and selected scATAC-seq profiles with a promoter ratio between 0.2 and 0.6 and a log10-transformed unique molecular identifier count [log10(UMI)] between 3 and 5. This process resulted in 4,100 cells selected from the 10X mouse brain dataset and 15,136 cells selected from the MOs-M1 dataset. Using the same filtering criteria to remove low-quality cells, we selected 9,646 cells from the MOs-M1 dataset for further analysis.

We then performed clustering on the lower-dimension representation learned by SAILER with a Louvain algorithm on a KNN graph. We applied t-SNE to generate a 2D visualization of the landscape. As cell labels are not available, we next visualized the activity scores of several marker genes to justify the clustering results. We selected several marker genes from the gene annotation file to obtain gene read counts within each cell. To avoid extreme sparsity and discontinued values, we adopted MAGIC to smooth the gene-cell matrix to obtain the final gene-level expression matrix. For each cell and each marker gene of interest, we applied gene expression values corresponding to each cell and denoted them by color in the t-SNE plot.

## 3 Results

We applied SAILER on both simulated and real datasets and carried out comprehensive performance benchmarking with existing methods, as discussed in the following sections below.

### 3.1 Extensive cell-to-cell variations in scATAC-seq data arise from confounding factors rather than biological heterogeneity

We found that, in addition to the underlying cell states, confounding factors from various sources significantly contribute to the cellular heterogeneity in scATAC-seq experiments. For instance, we extracted two mouse brain scATAC-seq datasets – one from the 10X genomics website (10X) and one from the SnapATAC website (MOs-M1) (see details in the Methods section). We uniformly processed these two datasets and found that the number of fragments within the same dataset varied significantly. For example, the uniquely mapped read counts per cell ranged from 1,500 to 6,000 for the MOs-M1 dataset (**Fig. 2**). Moreover, datasets generated from different labs showed distinct signatures. Specifically, the MOs-M1 dataset sample had fewer reads per cell but was highly enriched in promoter regions (median read count 3.506 vs. 4.236, promoter ratio 0.337 vs. 0.290). Most existing methods ignore such confounding factors, resulting in biased latent cell representations in dimensional reduction.

**Fig. 2.**
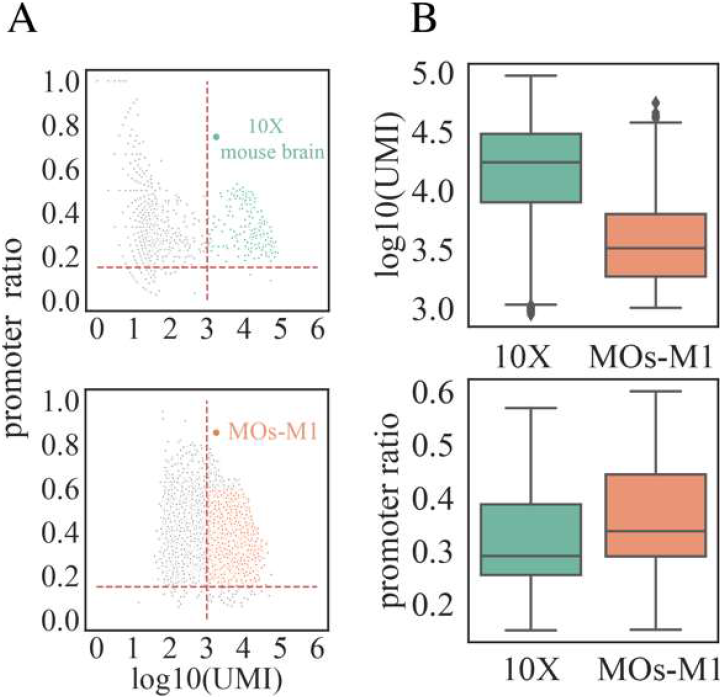
Visualization of confounding factors. **(A)** Scatter plots of a 10X mouse brain dataset (10X) and a mouse secondary cortex MOs-M1 dataset (MOs-M1). For all the cells in each dataset, we kept those with log10(UMI) between 0.3 and 0.5 and promoter ratio between 0.2 and 0.6. **(B)** Boxplots of read depth and promoter ratio comparison between selected cells from each dataset.

### 3.2 SAILER learns robust latent cell representations invariant to various confounding factors in simulated data

Here, we extensively benchmarked SAILER with existing methods using simulated data representing various application scenarios.

First, we simulated a deeply sequenced scATAC-seq dataset from five cell types, with varying mapping reads per cell. We learned the latent cell representations using SAILER, SCALE, and LSI as the input for the same clustering process. As shown in **Fig. 3A**, linear methods like LSI could not capture the complex dependencies among the peaks and hence failed to distinguish the rare cell type from the major cell types (red dots in the gray cluster). In contrast, both SAILER and SCALE used a nonlinear dimension reduction via fully connected neural networks and were able to report five clearly separable clusters. Furthermore, LSI and SCALE have a limited or no explicit module for correcting read depth effects. As a result, their L-shaped cell clusters are severely confounded by fragment counts, as reflected by the smooth transition from shallowly sequenced cells to densely sequenced ones within each cluster (the yellow to red pattern in **Fig. 3A**, Sim1). Such artifacts would be further amplified in the downstream imputation analysis, because cells with more mapped reads will exhibit even larger read counts after incorporating information from their similarly deeply sequenced neighbors. On the contrary, SAILER penalizes such depth effects by introducing an extra penalty term to force the latent cell representations to be as independent as possible to fragment counts per cell, resulting in compact round-shaped clusters with almost random read count distributions **(Fig. 3A**, Sim1). This observation is consistent with the quantitative measure of the mutual information ***I***(***z***, ***c***) between read counts and cell embeddings, where SAILER reported the lowest ***I***(***z***, ***c***) at 0.107 among all three methods (0.290 and 0.610 for SCALE and LSI, respectively, **Table 1**, Sim1). Thus, SAILER effectively removes confounding factors and learns robust cell representations.

**Fig. 3.**
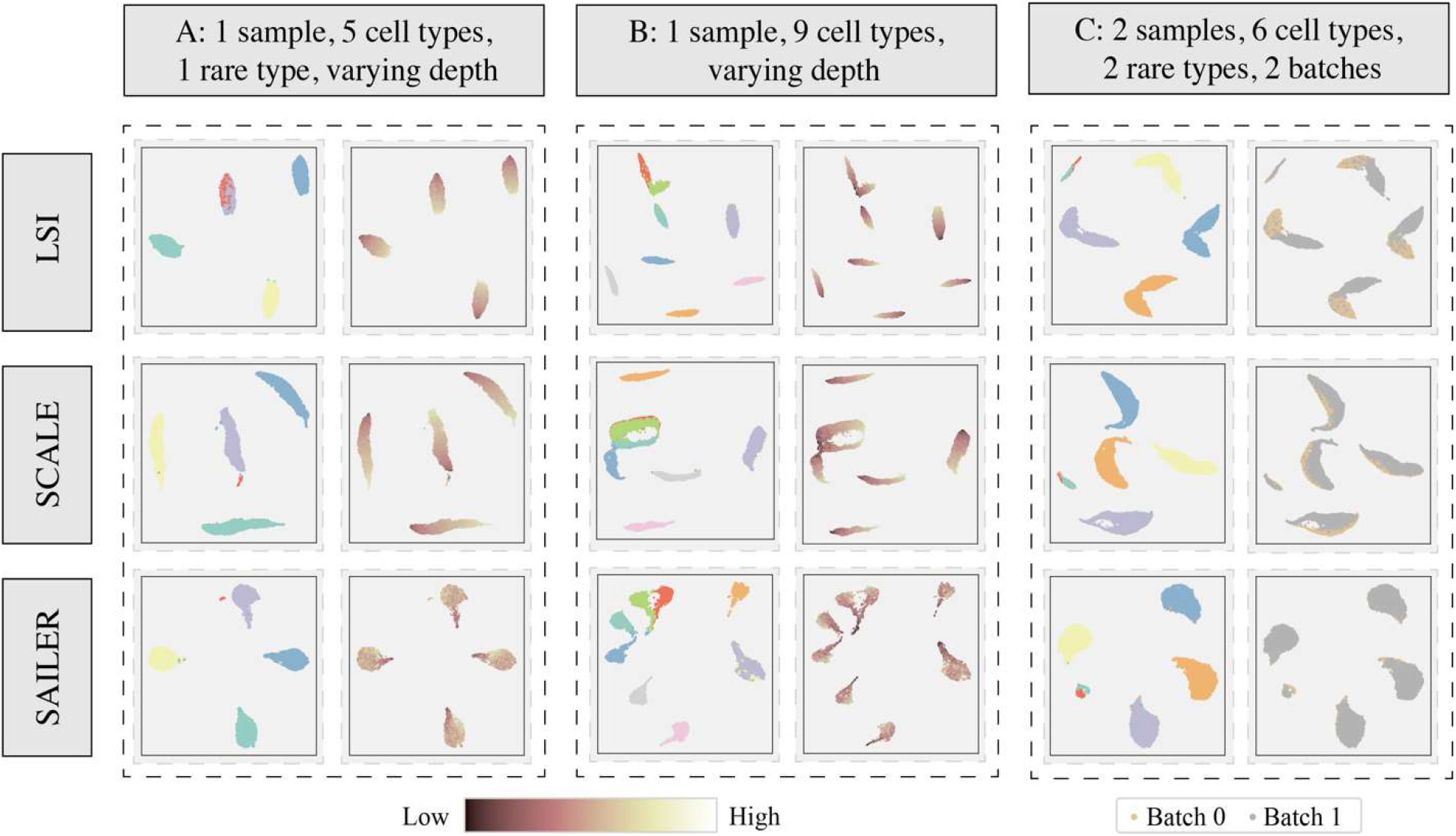
Results on simulation datasets. **(A)** 2D visualization of learned latent representations of LSI (top), SCALE (middle), and SAILER (bottom) on the Sim1 dataset. The left column shows the distribution of cell types. The right column shows the distribution of read depth indicated by color depth. **(B)** 2D visualization of learned latent representations of LSI (top), SCALE (middle), and SAILER (bottom) on the Sim2 dataset. The left column shows the distribution of cell types. The right column shows the distribution of read depth indicated by color depth. **(C)** 2D visualization of learned latent representations of LSI (top), SCALE (middle), and SAILER (bottom) on the Sim3 dataset. The left column shows the distribution of cell types. The right column shows the distribution of cells from different batches.

**Table 1.**
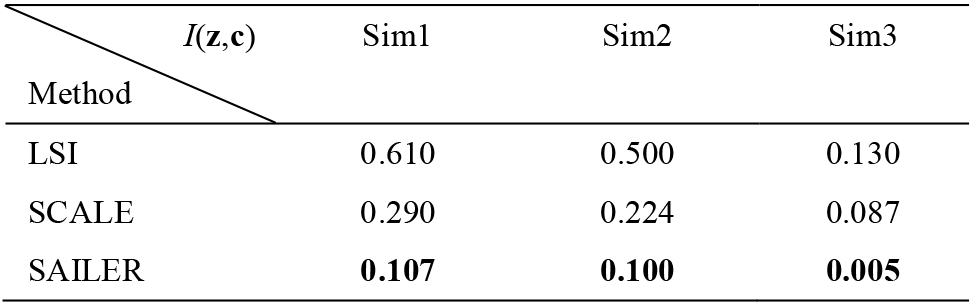
Mutual Information between the latent representation and confounding factors on simulation datasets.

We further simulated another shallowly sequenced dataset with fewer fragments per cell but more cell types, in order to conduct clustering performance benchmarking under more complicated (and realistic) scenarios. As shown in **Fig. 3B**, SCALE and LSI failed to separate two major cell types by reporting completely overlapped clusters (yellow and purple dots in **Fig. 3B**). Similar to the previous simulation, we observed clear low-to-high read count transitions within their reported clustering, indicating severe read depth artifacts. By contrast, SAILER distinguished cell types from distinct cell states into clear groups and demonstrated homogeneous read counts within each cluster (bottom row, **Fig. 3B**), indicating effective read depth bias removal. As expected, SAILER also showed the smallest amount of mutual information between fragment counts and latent cell representations (0.100 vs. 0.224 for SCALE and 0.500 for LSI, **Table 1**, Sim2), confirming the efficacy of its invariant representation learning scheme.

Lastly, we designed a third simulation dataset to mimic the scATAC-seq integration scenario with obvious batch effects for all three methods. We used latent representations to generate 2D visualizations with UMAP, as shown in **Fig. 3C**. We applied both batch information (right column) and cell-type information (left column) to annotate the plots. As shown in the right column, even though LSI and SCALE can marginally cluster the same type of cells, there are still clear boundaries between these batches. However, SAILER merges different batches very well, indicating that this method can remove batch information and retrieve the true distribution of cell biological states via the invariant latent representations. In order to quantitively measure how well these two batches are merged using different methods, we also calculated the mutual information between the batch information and each dimension of the latent representations (i.e., *I*(***z***, ***c***)), as shown in **Table 1**. SAILER still had the lowest value of mutual information (0.005, compared to 0.130 and 0.087). Note this dataset contains two sample-specific rare cell types (red and green dots, **Fig. 3C**), representing a potentially common situation in which certain rare cell types only appear in a few batches. LSI and SCALE completely merged the rare cell types together; however, SAILER was able to distinguish these two cell types after removing depth variation and batch effects from the latent representation.

### 3.3 SAILER outperforms existing methods in atlas-scale data analysis by reporting clearly separable clusters

To test the efficiency and accuracy of SAILER in a large-scale analysis, we benchmarked our method on a mouse atlas scATAC-seq dataset with ~80k cells from 40 cell types with substantial read depth variations, as shown in **Fig. 4**. We benchmarked SAILER with the GMM VAE in SCALE, and SnapATAC, the leading and only algorithm that was able to perform large-scale scATAC-seq analysis in a previous benchmarking study (Chen *et al*., 2019). As shown in **Fig. 4**, SAILER can learn robust cell representations that generate tight and clearly separable clusters, as compared to other methods.

**Fig. 4.**
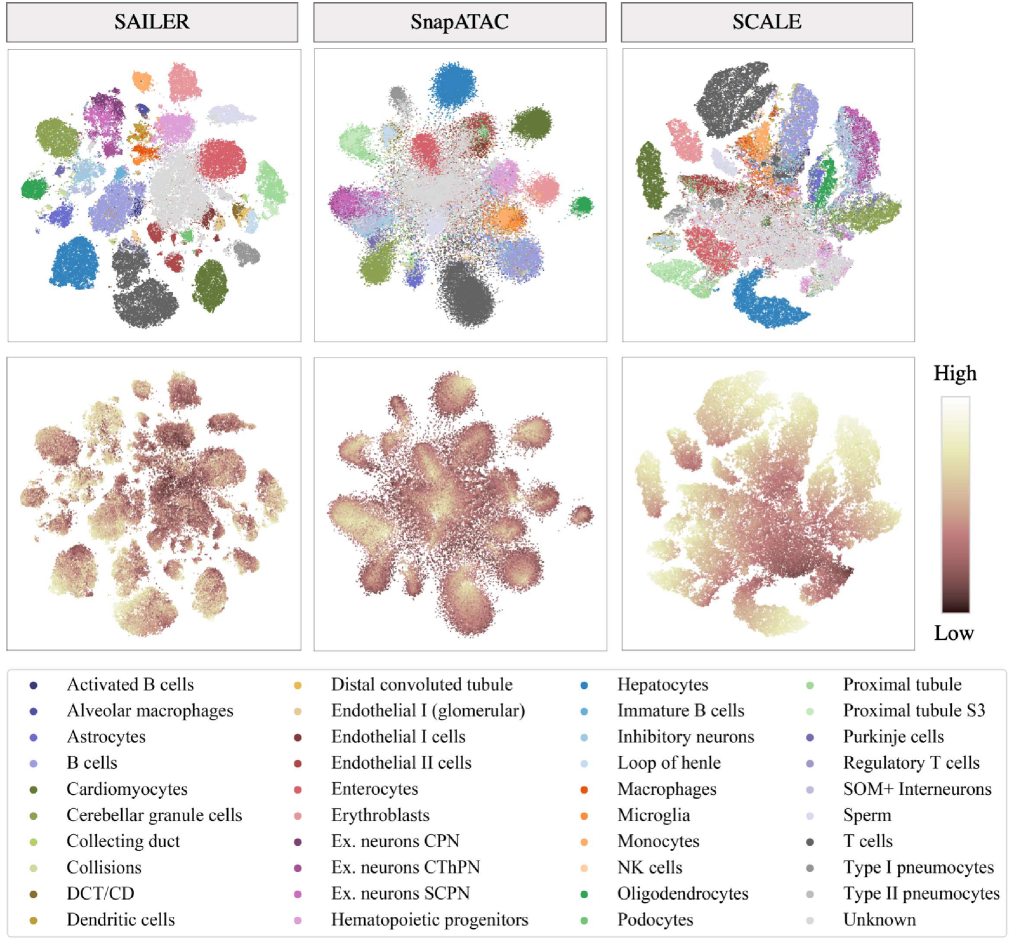
Results on the mouse atlas dataset. t-SNE visualization of lower-dimensional representation generated by SAILER (left), Snap-ATAC (middle), and SCALE (right). The first row shows the distribution of cell types. The second row shows the distribution of read depth indicated by color depth.

Besides, due to the lack of effective read depth removal, clustering results from SCALE are significantly confounded by the total number of fragments per cell. Specifically, the direct neighbors of deeply sequenced cells in SCALE’s reports are mostly those with higher read counts in each cluster (light dots in the bottom line, **Fig. 4**). This read depth effect will severely impact the subsequent imputation analysis, as depth imbalance among cells will be amplified when considering the neighbors. SnapATAC tends to remove such depth effects by regressing out fragment counts per cell in the cell-to-cell similarity calculation. As a result, its identified clusters are less affected by read depth. However, several internal groups were mixed together without clear separation, probably due to its independence and the equal contribution assumption among various genomic regions in the Jaccard distance calculations. Unlike SnapATAC, which requires a separate process for depth variation removal, SAILER integrates depth removal into the learning process – the fully connected neural network layers in SAILER allow nonlinear interactions among different genomic regions to better separate cells from different biological states, while the extra mutual information penalty term effectively removes read depth effects. This unified framework of SAILER makes each task aware of the other tasks, resulting in noticeably improved clustering results. This noticeable improvement can also be seen in the resulting NMI and ARI scores (**Table 2**). For instance, SCALE and SnapATAC reported NMI scores of 0.557 and 0.748, respectively, using known cell type-level labels, whereas SAILER showed a significantly higher NMI of 0.799. Moreover, SAILER reported lower mutual information (0.04), compared with 0.127 in SnapATAC and 0.279 in SCALE, suggesting successful depth effect removal for this method.

**Table 2.** Evaluation results on the mouse atlas dataset

| Method | ARI | NMI | $I(\mathbf{z}, \mathbf{c})$ |
| --- | --- | --- | --- |
| SAILER | **0.575** | **0.799** | **0.040** |
| SnapATAC | 0.538 | 0.748 | 0.127 |
| SCALE | 0.315 | 0.557 | 0.279 |

It is worth mentioning that the complexity of the batch-based training process increases linearly with the size of the input dataset, resulting in better scalability of SAILER to efficiently process millions of cells in multi-sample analyses. However, the polynomial regression approach used in SnapATAC increases quadratically as the number of cells increases. Chen *et al*. reported that Snap-ATAC takes nearly 12 hours to process the entire mouse atlas dataset (Chen *et al*., 2019), while SAILER can complete this process within 6 hours trained for 400 epochs. This further demonstrates the advantage of the deep learning method when scaling to very large datasets.

Moreover, we also followed the preprocessing procedures for subsampling by 10k cells for performance benchmarking with 17 other methods, as most methods cannot handle an atlas-scale dataset. Instead of cell-type labels, we used the same tissue-level cell labels for comprehensive clustering benchmarking. When applied to the subsampled dataset, SAILER still achieved the highest ARI (0.397) among all methods (with the 17 other methods ranging from 0.009 to 0.363). This further demonstrates the effectiveness of our method.

### 3.4 SAILER can effectively remove batch effects in multi-sample scATAC-seq integration

Another common source of confounding factors are batch effects in multi-sample scATAC-seq analysis, where samples may be processed and sequenced from different labs or even sequencing platforms with distinct sample-specific signatures. To evaluate the performance of our method in such scenarios, we applied SAILER on two mouse brain scATAC-seq samples from two sources – one mouse brain dataset from the 10x Genomics website (10X) and one generated from mouse secondary cortex brains (Fang *et al*., 2019).

For fair performance benchmarking, we uniformly processed these two datasets to identify cells from random barcodes using the default parameters in SnapATAC (Fang *et al*., 2019). Specifically, after removing barcodes with less than 1,000 fragments and keeping the remaining ones with promoter ratios between 0.2 and 0.6, we identified 4,100 and 9,646 cells from these two samples (see details in the Methods section). Starting from the same tissue, we found that these two samples generated from different labs showed distinct fragment signatures. For instance, the dataset from the 10X Genomics website demonstrated a higher mean read coverage per cell (log(UMI) = 4.149 vs. 3.547, P-value = 10e-15 using the two-sided Wilcoxon test) and a lower mean promoter ratio (0.320 vs. 0.367, P-value = 2.48e-87 using the two-sided Wilcoxon test). After pre-processing, we projected the remaining cells into a ten-dimensional space using SAILER and SCALE, and then generated a KNN graph (k=16) and performed clustering via the Louvain algorithm. We also used t-SNE to map the ten-dimensional cell representations onto a 2D space for visualization and labeled the sample IDs using different colors in **Fig. 5**. In the ideal case, a good computational method should overcome batch effects by reporting cell clusters with homogenous sample ID distributions. However, due to the lack of an appropriate batch effect removal module, we found that clusters reported by SCALE were predominantly driven by sample effects rather than the true biological states of the cells (**Fig. 5A**). In contrast, SAILER effectively removed batch effects by introducing an additional penalty to reduce the mutual information *I*(***z***, ***c***) between the variant component and the batch component in the objective function. As a result, the different samples were homogeneously mingled in the clearly separated clusters reported by SAILER (yellow and grey dots in **Fig. 5A**).

**Fig. 5.**
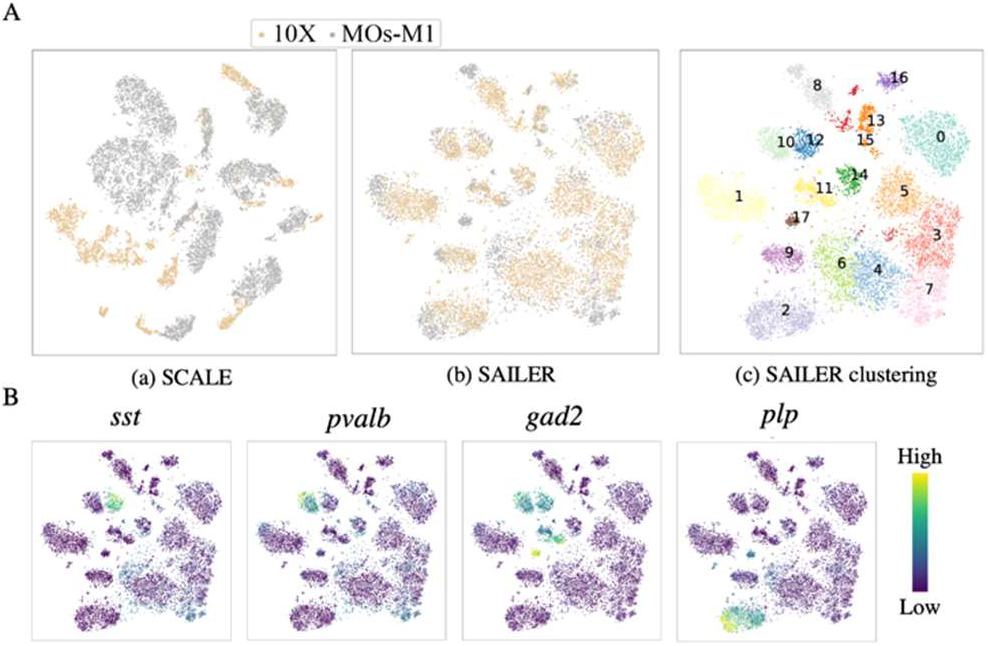
Results on mixed mouse brain datasets. **(A)** Clustering result comparison of SCALE and SAILER on two batches of mouse brain cell samples. Clustering result (a) using SCALE, (b) using SAILER, and (c) using SAILER but colored and labeled with numbers calculated using the Louvain method based on the KNN graph. **(B)** Clustering result of SAILER on two batches of datasets but colored with four marker gene scores, namely *sst, pvalb, gad2,* and *plp1*. The brighter the color, the higher the gene score shown for those cells.

To test whether these SAILER-reported clusters represent distinct biological cell states, we calculated the overall chromatin accessibility scores of well-known marker genes (Fang *et al*., 2019) and labeled cells using the activity scores of the marker genes. As shown in Fig. 5B, SAILER identified clearly separable cell clusters that correspond well with the activities of the marker genes (*sst, pvalb, gad2,* and *plp1*). For instance, sst is a well-known marker gene widely expressed in inhibitory neurons. SAILER homogeneously grouped together sst-enriched cells from different batches, demonstrating its ability to appropriately remove batch effects while retaining the true cell-cell variability.

### 3.5 SAILER can precisely reconstruct a chromatin accessibility landscape free of various confounding factors

Despite high throughput in revealing epigenetic heterogeneity, scATAC-seq experiments suffer from severe missingness by reporting only a few thousand fragments in the entire genome. Therefore, accurate chromatin landscape reconstruction and imputation are essential to uncovering the full regulatory potential within a cell. However, very few computational methods are designed explicitly for chromatin accessibility imputation.

Here, we took advantage of the deep generative model and its invariant representation to reconstruct a full chromatin accessibility landscape that is independent of sequencing depth and batch effects. During imputation, we fixed the values of the confounding variables, such that the variations of the reconstructed scATAC-seq data only depend on the invariant representation ***z***, which reflects the intrinsic variation of biological states.

To further demonstrate this, we performed imputation on the third simulation dataset (Sim3) with two simulated samples. SCALE is currently the only available method designated for imputing scATAC-seq data. LSI has no direct imputation module, we added MAGIC as suggested for benchmarking (Granja *et al*., 2020). First, SAILER, MAGIC, and SCALE generated the imputed data. These data, along with the raw data, were then processed by PCA and visualized with UMAP in 2D. From the PCA embeddings shown in **Fig. 6**, we found that the imputation data of SCALE were severely affected by depth variation and batch effects. We observed similar results with MAGIC, where after imputation, the same types of cells from different batches were divided into separate clusters in the PCA embedding. However, the imputed data by SAILER did not show separate clusters from different batches. Moreover, the rare cell types (shown in green and red, **Fig. 6**) were separable in the PCA embedding, which was not the case for SCALE or MAGIC. The results indicate that, without proper removal of confounding factors during imputation, the imputed data show clear variations that correlate with confounding factors. In addition, the data diffusion strategy used in MAGIC is not friendly to rare cell types, as the rare cells can be easily overwhelmed by the major cell types. Thus, compared with SCALE and MAGIC, SAILER is the only method capable of removing confounding factors from imputation data, while preserving unique information from rare cell types.

**Fig. 6.**
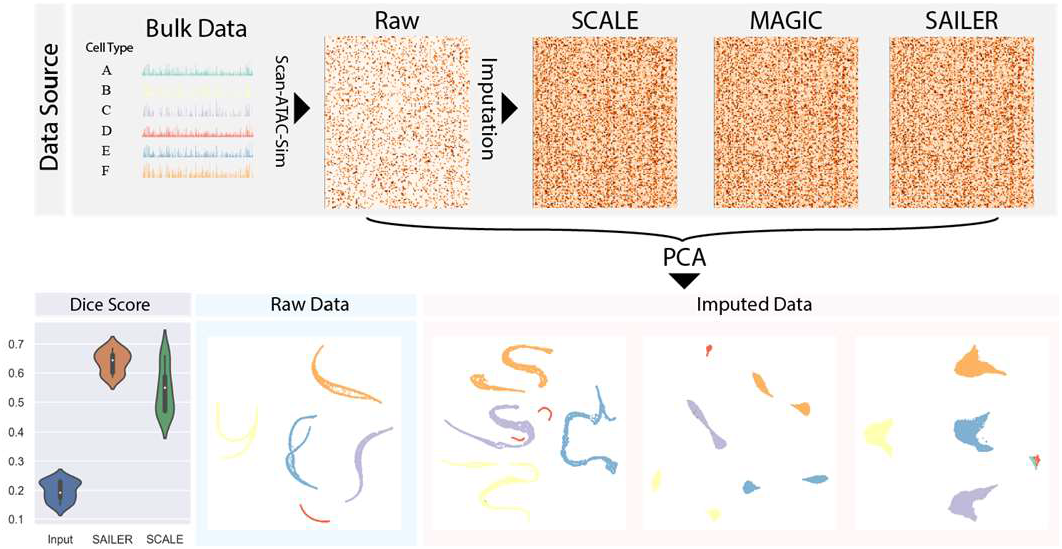
Imputation pipeline and results. Simulated data (Sim3) with 2 batches is generated by Scan-ATAC-Sim tool. Imputed data is generated by running SCALE, MAGIC, and SAILER, respectively. Imputed data is then subject to PCA and visualized by UMAP. Dice Score is computed between each imputed data and the Bulk Data. The Dice score between Input data and Bulk data is also shown as baseline.

As the bulk ATAC-seq data used to simulate the single-cell data is available, we used the bulk data as the ground truth and calculated the DSC for each imputation method. The DSC (also known as the F1 Score) is a harmonic mean of the precision and recall. Because scATAC-seq is imbalanced in 0/1 entries, we used DSC as a balanced metric to evaluate the imputation performance. We generated a violin plot to show the DSC distributions of raw single-cell data, SAILER, and SCALE. As shown in **Fig. 6**, SAILER and SCALE both achieved higher DSC scores compared to the raw data, indicating that both methods generate reasonable imputation results. SAILER achieved a higher mean DSC compared with SCALE (0.64 vs. 0.54), further demonstrating the effectiveness of invariant representation learning.

## 4 Discussion

In this work, we developed a scalable and accurate single-cell ATAC-seq processing and integration method called SAILER via efficient invariant representation learning. As compared with previous methods, SAILER has **three distinct characteristics** designed explicitly for single-cell data analysis – 1) it utilizes nonlinear dimension reduction via fully connected neural networks in a deep generative framework to handle complex dependencies among various peaks; 2) it dissociates cell-state-related biological variations from those arising from confounding factors (e.g., read depth and batch effects) to faithfully embed the cells into a low-dimensional latent space to facilitate various downstream analyses, such as cell clustering and imputation; 3) it is easily scalable to large-scale single-cell data analysis accelerated using GPU parallelism.

We applied SAILER to various simulated and real scATAC-seq datasets and comprehensively compared its performance with *state-of-the-art* analysis pipelines. We showed that SAILER’s robust cell embeddings can effectively remove noise impacts from different sources and improve clustering and imputation results on all of the benchmark datasets. We should note that the invariant representation learning framework presented here is general and can be applied to other types of high-throughput genomic data like scRNA-seq and single-cell DNA methylation, or to joint analysis of multi-modality single-cell genomics data. Specifically, several single-cell multi-omics technologies have recently emerged for measuring multiple types of molecules in the same cell (Jin *et al*., 2020). To achieve this, we could apply a multi-modal VAE to encode a variational posterior jointly from single-cell multimodal omics inputs using deep neural networks, where the resultant latent space factors into a shared subspace to profile cell states or functions for individual cells and private subspaces could be used to solve specific technical issues for each modality.

In summary, we developed a deep generative model, SAILER, for learning robust latent cell representations invariant to changes in various noise factors, which has not been possible with most current scATAC-seq analysis tools. Given the fast-expanding collection of publicly available single-cell sequencing data, we envision that the SAILER framework can serve as a powerful tool to remove impacts from confounding factors and uncover cellular heterogeneity across diverse cell states and conditions in large-scale single-cell omics data analysis.

## Acknowledgements

We thank A Hwang, L Zhang, and members of Xie lab for helpful discussions.

